# A comprehensive examination of ACE-2 receptor and prediction of spike glycoprotein and ACE-2 interaction based on in silico analysis of ACE-2 receptor

**DOI:** 10.1101/2021.12.14.472331

**Authors:** Nehir Özdemir Özgentürk, Emre Aktaş

## Abstract

ACE-2 receptor plays a vital role not only in the SARS-CoV-induced epidemic but also in some diseases. Studies have been carried out on the interactions of ACE-2-SARS-CoV proteins. However, comprehensive research has not been conducted on ACE2 protein by using bioinformatic tools. The present study especially two places, G104 and L108 points, which are effective in protecting the structure of the ACE-2 protein, play a critical role in the biological functioning of this protein, and play an essential role in determining the chemical-physical properties of this protein, and play a crucial role for ACE-2 protein-SARS CoV surface glycoprotein, were determined. It was also found that the G104 and L108 regions were more prone to possible mutations or deletions than the other ACE-2 protein regions. Moreover, it was determined that all possible mutations or deletions in these regions affect the chemical-physical properties, biological functions, and structure of the ACE-2 protein. Having a negative GRAVY value, one transmembrane helix, a significant molecular weight, a long-estimated half-life as well as most having unstable are results of G104 and L108 points mutations or deletions. Finally, it was determined that LQQNGSSVLS, which belong to the ACE-2 protein, may play an active role in binding the spike protein of SARS-CoV. All possible docking score results were estimated. It is thought that this study will bring a different perspective to ACE-2 _SARS-CoV interaction and other diseases in which ACE-2 plays an important role and will also be an essential resource for studies on ACE-2 protein.

## Introduction

Over one year, severe acute respiratory syndrome coronavirus 2 (SARS-CoV-2) has caused a pneumonia outbreak in Wuhan city, China, resulting in a global spread(Wu et al.,2020). SARS-COV-2 has spike proteins that have two functional subunits (S1, S2). The S1 subunit plays a role in binding to the ACE-2 (Angiotensin-converting enzyme 2) receptor on the host cell, which is crucial for the virus entry into epithelial cells (Chan-Yeung M and Yu., 2003; Belouzard, S. et al.,2009; Hui, D. S. et al.,2020; Wu et al.,2020). Therefore, studies on the ACE-2 protein have increased after this outbreak. Nevertheless, ACE-2 is not only a receptor that causes coronavirus to enter the cell, but also Ang II (vasoconstrictor angiotensin II) has been determined to act as an essential negative regulator of RAS (renin-angiotensin system). - inflammation, anti-hypertension, anti-thrombus, and anti-fibrosis activity have been found to play several roles in improving cardiovascular protection(Xiaocong et al.,2020; Hoffmann et al.,2020;Aktas et al.,2021;Aktas E.,2021). It has also been noted that ACE2 plays a vital role in acute respiratory distress syndrome (ARDS) and acute lung injury (ALI) induced by the deadly avian influenza A (H5N1, H7N9)(Zou et al.,2014; Yang et al.,2014). Diminazene aceturate (DIZE), the anti-trypanosomal agent, has been reported as an ACE2 activator with a structure similar to the ACE2 activator. The same study suggested that DIZE exerts protective effects in cardiovascular disease by modulating ACE2 activation and expression to increase Ang 1-7 production and improve vascular function(Qaradakhi et al.,2020). Pang et al. Stated in their study that adding different numbers of rhACE2 to patients affected by the virus could be a potential treatment for SARS-CoV-2 infection(Xiaocong et al.,2020) ACE2 expression is severely decreased in patients with pulmonary fibrosis(Li et al.,2008). Therefore, recombinant human ACE2 (rhACE2) injection is considered to treat ARDS and pulmonary arterial hypertension (Zhang and Baker,2017). When new recombinant vaccine designs, reverse vaccinology (RV) in silico approach delivers comprehensive initial prediction about vaccine nominee using relevant sequence. Using the RV in silico approach is crucial because it predicts the antigenicity, the epitope regions of B and T cells, and other parameters such as a signal peptide, physicochemical parameters, and solubility about targeted proteins (Rashid et al.,2019;Can et al.,2020). Bioinformatics has recently played a pivotal role in the immune system, predicted possible mutations in proteins, and predicted protein structure. Result acquired from in silico prediction has extreme importance for preventing the failures encountered at the end of wet-lab studies or even late stages of clinical trials(Dangi et al.,2018;Rashid et al.,2019;Can et al.,2020).

Understanding the structure of ACE-2 protein is essential(Chan-Yeung M and Yu., 2003; Li et al.,2008; Belouzard, S. et al.,2009; Hui, D. S. et al.,2020; Wu et al.,2020; Qaradakhi et al.,2020; Hoffmann et al.,2020;Aktas et al.,2021;Aktas E.,2021). This study aimed to investigate the structure of ACE-2(NCBI Reference Sequence: NP_068576.1) in depth using bioinformatics tools. The results were obtained in which regions of ACE-2 were not unregulated/unstructured regions. It was predicted that especially G104 and L108 sites could be affected by all possible point mutations and deletions. Prediction of whether the substitution or deletion of amino acids(G104, L108) affects the biological function of the ACE-2 receptor and prediction of protein stability changes upon single-site mutations results predicted were analyzed. These results’ physical and chemical properties were found with bioinformatics tools, and all results were predicted to support each other. It was predicted that these G104 and L108 regions crucial for not only Spike-ACE-2 protein interaction but also ACE-2’s structural, chemical, and physical properties. It is thought that all these studies can give a new perspective to some of ACE-2’s structural, chemical, and physical properties. Finally, all these preliminary studies will be an essential resource for studies on ACE-2 receptors in the future.

## Method

### ACE□2 isolate, and it’s variant protein

NCBI (National Center for Biotechnology Information) (https://www.ncbi.nlm.nih.gov) was used to obtain the whole sequence of ACE-2 protein (Accession number: NP_068576.1 and sequenced in January 2021), and Molecular Evolutionary Genetic Analysis(MEGA) was used to alignment(Kumar et al.,2016).

### Prediction unregulated / unstructured regions for ACE-2 protein

DisEMBL was used for the prediction of disordered/unstructured regions within the ACE-2 protein sequence. As no clear definition of disorder exists, it has been developed parameters based on several alternative definitions (Disordered by Loops/coils definition, Disordered by Hot-loops definition, and Disordered by Remark-465 definition), and introduced a new one based on the concept of “hot loops,” i.e., coils with high-temperature factors(Linding et al.,2003).

### Predicting possible change, and its effect on the biological function of the ACE-2 protein

PROVEAN (Protein Variation Effect Analyzer) was used to predict whether an amino acid substitution or indel impacts the biological function of a protein(Choi et al.,2012).

### Prediction of ACE-2 protein stability changes

I-Mutant2.0 was used for the prediction of protein stability changes upon single-site mutations. I-Mutant2.0 evaluates the stability change upon single-site mutation starting from the protein structure or the protein sequence (Emidio et al., 2005).

### Prediction of wild-type and mutant-type ACE-2’ amino acid properties

HOPE was used to analyze the structural effects of a point mutation in a protein sequence. HOPE collected and combined available information from a series of web services and databases and produced a report, complete with results(Hanka et al.,2010).

### Prediction of physico-chemical parameters of ACE-2 protein

The reference genome proteins were investigated using Expasy ProtParam online server (https://web.expasy.org/protparam/) to predict physico-chemical properties(Gasteiger et al.,2005). The prediction of solubility was performed by SolPro (http://scratch.proteomics.ics.uci.edu/)(Cheng et al.,2005).

### Prediction evolutionary pattern of amino acids on ACE-2 protein

ConSurf was used to analyze the evolutionary pattern of the amino acids of the ACE-2 protein to reveal regions that are important for structure and function(Ashkenazy et al.,2016).

### Small tertiary structure predicted, and structures visualized

The tertiary structure of peptide was predicted by 3Dpro(Gaëlle et al.,2009), and all the structures are visualized using Chimera 1.15(Pettersen et al.,2004).

### Prediction of docking scores

HDOCK SERVER was used to predict the docking score between spike protein and ACE-2 receptor interaction(Yan et al.,2020).

## Results

The DisEMBL computational tool was used to predict the unregulated/unstructured regions of ACE-2’s FASTA (NCBI Reference Sequence: NP_068576.1). The DisEMBL tool uses different alternatives when trying to detect disorders(Linding et al.,2003). These are Disordered by Loops/coils definition, Disordered by Hot-loops definition, and Disordered by Remark-465 definition(Linding et al.,2003). It was chosen the regions which are common to all of these different alternatives (Table 1)(Linding et al.,2003). These common places are highlighted in black (Table 1). They are GSSVLSE(104-110 on the NP_068576.1) and DRKKKNKARS(767-777 on NP_068576.1). The amino acids shown in the upper case letters indicate the disordered regions (Table 1).

**Table 1.** Common unregulated/unstructured regions of ACE2 results predicted by disEMBL tool. These common regions, both location, and sequences are highlighted in black.

| Tool for predicting | Predicted Locations | Common Predicted Results |
| --- | --- | --- |
| DisEMBL |  |  |
| 1. Disordered by Loops/coils definition | 1. **103-111**, 128-151, 194-223, 250-267, 280-295, 317-357, 387-400, 410-432, 464-472, 487-511, 533-547, 559-567, 575-588, 594-615, 627-640, 668-698, 714-740, **766-805** | **GSSVLSE(104-110)**<br>**DRKK KNKARS(767-777)** |
| 2. Disordered by Hot-loops definition | 2. **104-113**, 683-700, **762-805** |  |
| 3. Disordered by Remark-465 definition | 3. **101-110**, **767-777** |  |

The results(belongs to NP_068576.1) in Table 1 were used to predict whether the substitution or change of an amino acid in the PROVEAN (Protein Variation Effect Analyzer) tool affects the biological function of the protein(Choi et al.,2012). All of the amino acids in these GSSVLSE (104-110 on the NP_068576.1) and DRKKKNKARS (767-777 on NP_068576.1) regions were point mutated one by one, and the results are shown in Table 2. All possible mutations at points G104*X* (representing all amino acids except X: G) and L108*X*(representing all amino acids except X: L) were predicted to result in deleterious consequences above the biological function of ACE-2 protein. All these deleterious results are shown in Table 2. Variants with the result of the PROVEAN tool score less than or equal to - 2.5 are considered “deleterious” and variants with a score above −2.5 are considered “neutral”(Choi et al.,2012). All results of both the G104*X*(representing all amino acids except X: G) and L108*X* (representing all amino acids except X:L)points were predicted less than −2.5(Table 2). The least result G104L is −6.606, while the maximum result G104D is −2.955. Besides, the minor result for L108 is −6.815(L108G), while the maximum result is −2.955(L108V)(Table 2). In addition to the results of possible mutations, it was predicted that a possible deletion at both the G104 and L108 points would be deleterious on the biological function of ACE-2 protein. The results of L108del is −9.391, while the G104del result is −9.872. Deletions K769del, K770del, K771del, N772del, K773del, A774del, R775del, S776del, G777del were not predicted to deleterious, while the remaining deletions were predicted to be deleterious(Table 2). While the most harmless result was G777del(score:-0.343), the most harmful was predicted as G104del(score:-9.872). All predicted deleterious results which belongs to other position are highlighted in black on Table 2.

**Table 2.**
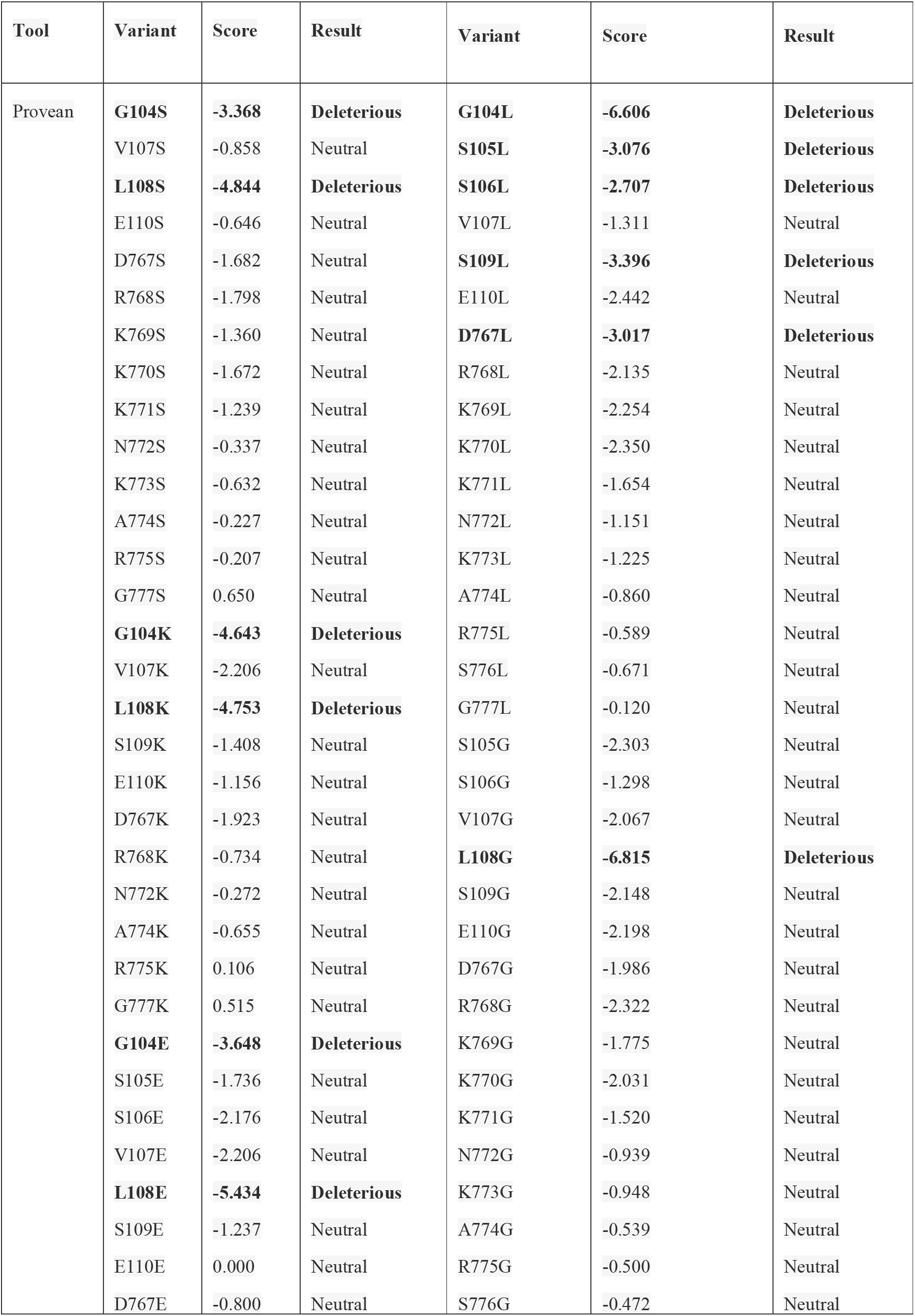
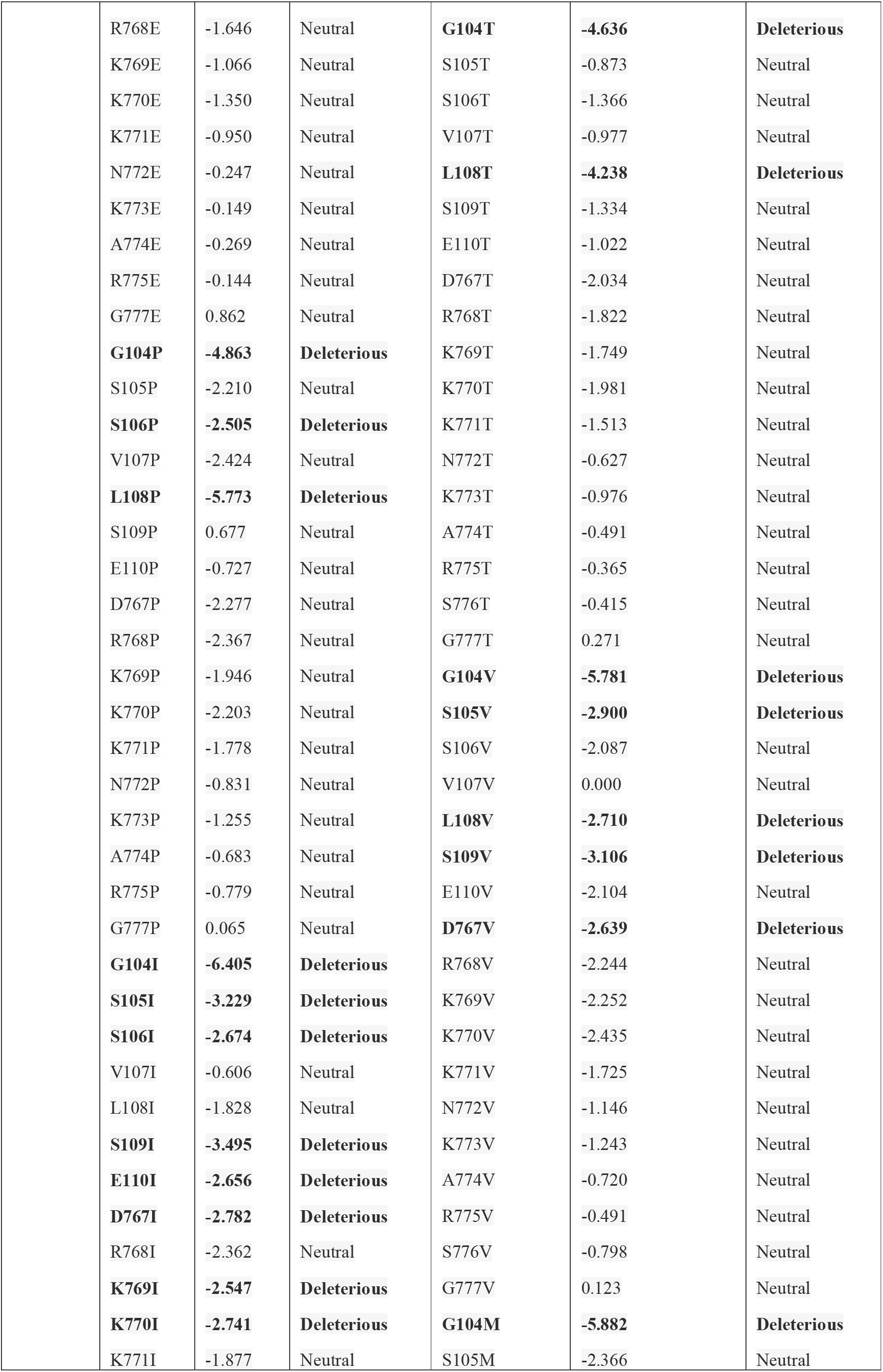
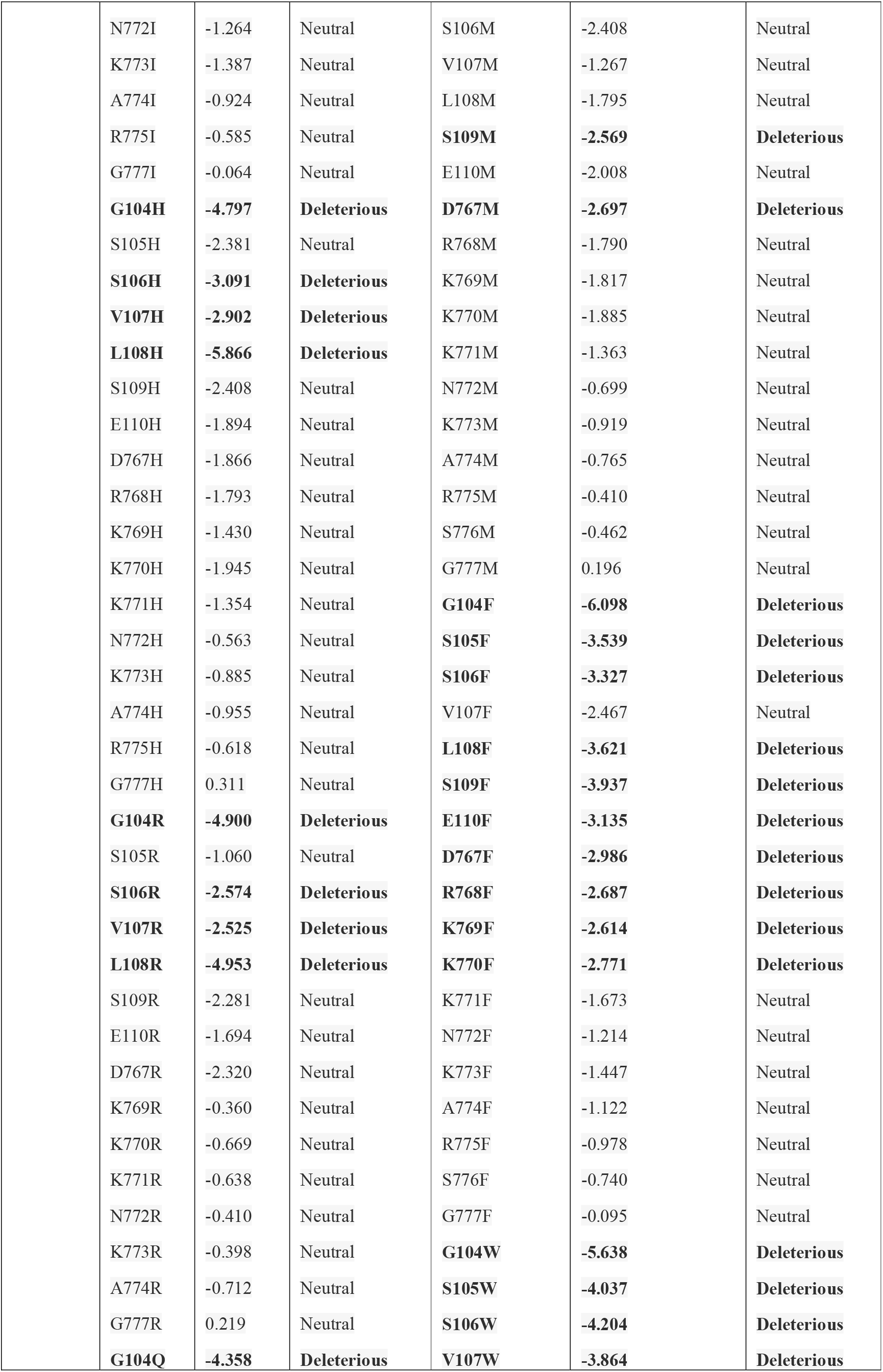
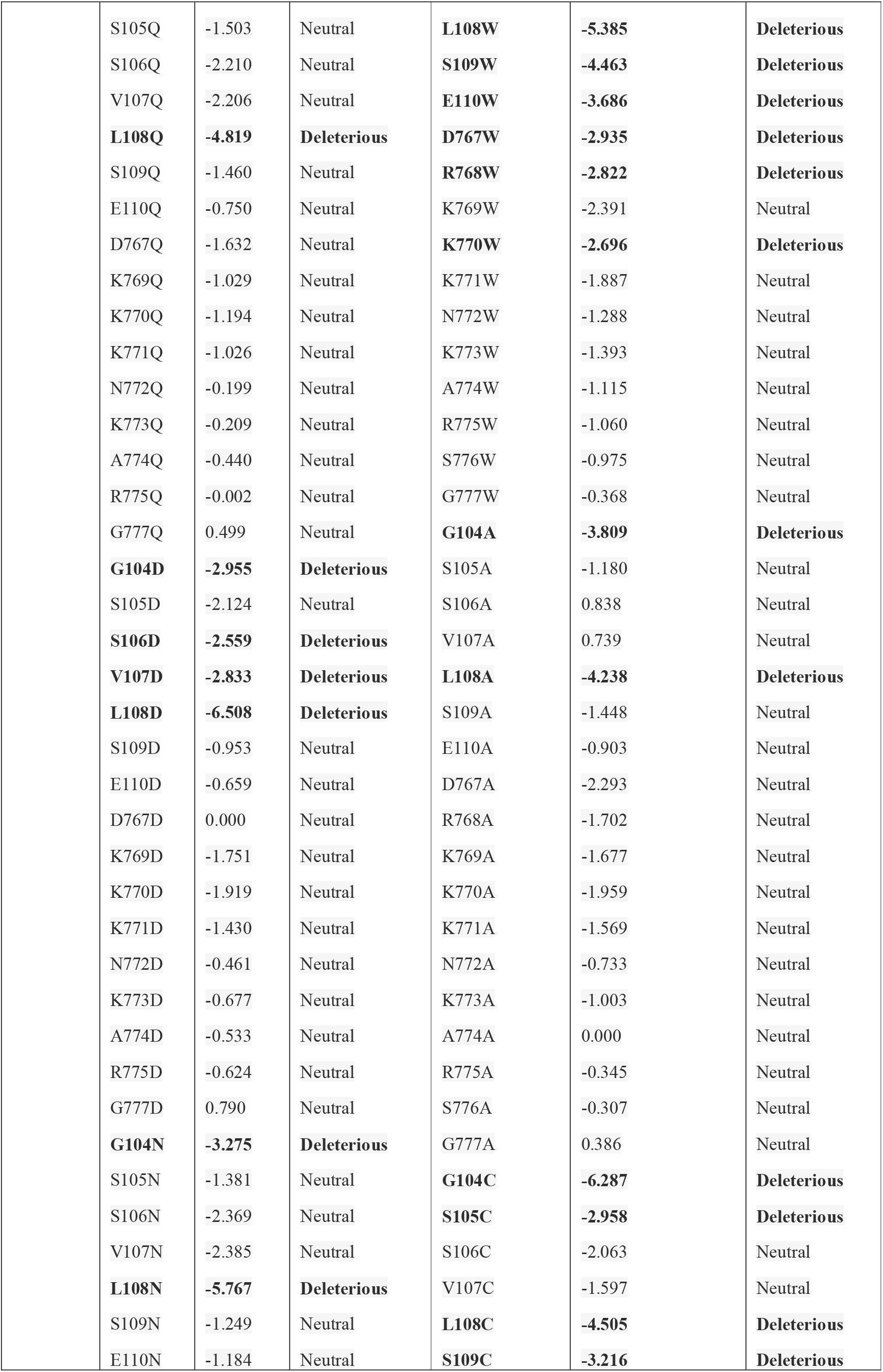
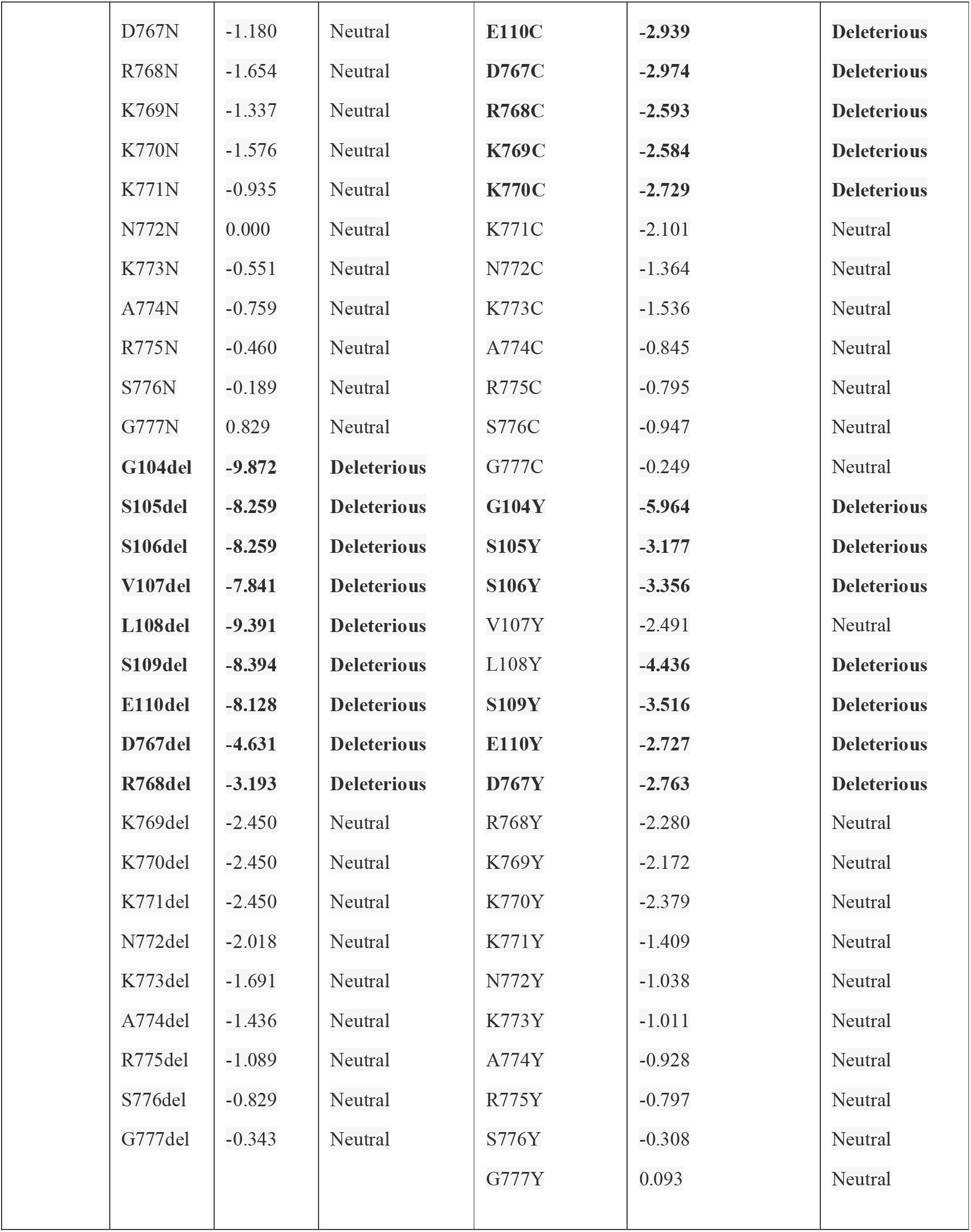
Prediction of whether the substitution or deletion of an amino acid affects the protein’s biological function (ACE-2:NP_068576.1) by PROVEAN tool. The GSSVLSE(104-110 on the NP_068576.1) and DRKKKNKARS(767-777 on NP_068576.1) are from Table 1. All predicted deleterious results and deletion results are highlighted in black.

| Tool | Variant | Score | Result | Variant | Score | Result |
| --- | --- | --- | --- | --- | --- | --- |
| Provean | **G104S** | **-3.368** | **Deleterious** | **G104L** | **-6.606** | **Deleterious** |
|  | V107S | -0.858 | Neutral | **S105L** | **-3.076** | **Deleterious** |
|  | **L108S** | **-4.844** | **Deleterious** | **S106L** | **-2.707** | **Deleterious** |
|  | E110S | -0.646 | Neutral | V107L | -1.311 | Neutral |
|  | D767S | -1.682 | Neutral | **S109L** | **-3.396** | **Deleterious** |
|  | R768S | -1.798 | Neutral | E110L | -2.442 | Neutral |
|  | K769S | -1.360 | Neutral | **D767L** | **-3.017** | **Deleterious** |
|  | K770S | -1.672 | Neutral | R768L | -2.135 | Neutral |
|  | K771S | -1.239 | Neutral | K769L | -2.254 | Neutral |
|  | N772S | -0.337 | Neutral | K770L | -2.350 | Neutral |
|  | K773S | -0.632 | Neutral | K771L | -1.654 | Neutral |
|  | A774S | -0.227 | Neutral | N772L | -1.151 | Neutral |
|  | R775S | -0.207 | Neutral | K773L | -1.225 | Neutral |
|  | G777S | 0.650 | Neutral | A774L | -0.860 | Neutral |
|  | **G104K** | **-4.643** | **Deleterious** | R775L | -0.589 | Neutral |
|  | V107K | -2.206 | Neutral | S776L | -0.671 | Neutral |
|  | **L108K** | **-4.753** | **Deleterious** | G777L | -0.120 | Neutral |
|  | S109K | -1.408 | Neutral | S105G | -2.303 | Neutral |
|  | E110K | -1.156 | Neutral | S106G | -1.298 | Neutral |
|  | D767K | -1.923 | Neutral | V107G | -2.067 | Neutral |
|  | R768K | -0.734 | Neutral | **L108G** | **-6.815** | **Deleterious** |
|  | N772K | -0.272 | Neutral | S109G | -2.148 | Neutral |
|  | A774K | -0.655 | Neutral | E110G | -2.198 | Neutral |
|  | R775K | 0.106 | Neutral | D767G | -1.986 | Neutral |
|  | G777K | 0.515 | Neutral | R768G | -2.322 | Neutral |
|  | **G104E** | **-3.648** | **Deleterious** | K769G | -1.775 | Neutral |
|  | S105E | -1.736 | Neutral | K770G | -2.031 | Neutral |
|  | S106E | -2.176 | Neutral | K771G | -1.520 | Neutral |
|  | V107E | -2.206 | Neutral | N772G | -0.939 | Neutral |
|  | **L108E** | **-5.434** | **Deleterious** | K773G | -0.948 | Neutral |
|  | S109E | -1.237 | Neutral | A774G | -0.539 | Neutral |
|  | E110E | 0.000 | Neutral | R775G | -0.500 | Neutral |
|  | D767E | -0.800 | Neutral | S776G | -0.472 | Neutral |

|  |  |  |  |  |  |
| --- | --- | --- | --- | --- | --- |
| R768E | -1.646 | Neutral | **G104T** | **-4.636** | **Deleterious** |
| K769E | -1.066 | Neutral | S105T | -0.873 | Neutral |
| K770E | -1.350 | Neutral | S106T | -1.366 | Neutral |
| K771E | -0.950 | Neutral | V107T | -0.977 | Neutral |
| N772E | -0.247 | Neutral | **L108T** | **-4.238** | **Deleterious** |
| K773E | -0.149 | Neutral | S109T | -1.334 | Neutral |
| A774E | -0.269 | Neutral | E110T | -1.022 | Neutral |
| R775E | -0.144 | Neutral | D767T | -2.034 | Neutral |
| G777E | 0.862 | Neutral | R768T | -1.822 | Neutral |
| **G104P** | **-4.863** | **Deleterious** | K769T | -1.749 | Neutral |
| S105P | -2.210 | Neutral | K770T | -1.981 | Neutral |
| **S106P** | **-2.505** | **Deleterious** | K771T | -1.513 | Neutral |
| V107P | -2.424 | Neutral | N772T | -0.627 | Neutral |
| **L108P** | **-5.773** | **Deleterious** | K773T | -0.976 | Neutral |
| S109P | 0.677 | Neutral | A774T | -0.491 | Neutral |
| E110P | -0.727 | Neutral | R775T | -0.365 | Neutral |
| D767P | -2.277 | Neutral | S776T | -0.415 | Neutral |
| R768P | -2.367 | Neutral | G777T | 0.271 | Neutral |
| K769P | -1.946 | Neutral | **G104V** | **-5.781** | **Deleterious** |
| K770P | -2.203 | Neutral | **S105V** | **-2.900** | **Deleterious** |
| K771P | -1.778 | Neutral | S106V | -2.087 | Neutral |
| N772P | -0.831 | Neutral | V107V | 0.000 | Neutral |
| K773P | -1.255 | Neutral | **L108V** | **-2.710** | **Deleterious** |
| A774P | -0.683 | Neutral | **S109V** | **-3.106** | **Deleterious** |
| R775P | -0.779 | Neutral | E110V | -2.104 | Neutral |
| G777P | 0.065 | Neutral | **D767V** | **-2.639** | **Deleterious** |
| **G104I** | **-6.405** | **Deleterious** | R768V | -2.244 | Neutral |
| **S105I** | **-3.229** | **Deleterious** | K769V | -2.252 | Neutral |
| **S106I** | **-2.674** | **Deleterious** | K770V | -2.435 | Neutral |
| V107I | -0.606 | Neutral | K771V | -1.725 | Neutral |
| L108I | -1.828 | Neutral | N772V | -1.146 | Neutral |
| **S109I** | **-3.495** | **Deleterious** | K773V | -1.243 | Neutral |
| **E110I** | **-2.656** | **Deleterious** | A774V | -0.720 | Neutral |
| **D767I** | **-2.782** | **Deleterious** | R775V | -0.491 | Neutral |
| R768I | -2.362 | Neutral | S776V | -0.798 | Neutral |
| **K769I** | **-2.547** | **Deleterious** | G777V | 0.123 | Neutral |
| **K770I** | **-2.741** | **Deleterious** | **G104M** | **-5.882** | **Deleterious** |
| K771I | -1.877 | Neutral | S105M | -2.366 | Neutral |
| N772I | -1.264 | Neutral | S106M | -2.408 | Neutral |
| K773I | -1.387 | Neutral | V107M | -1.267 | Neutral |
| A774I | -0.924 | Neutral | L108M | -1.795 | Neutral |
| R775I | -0.585 | Neutral | **S109M** | **-2.569** | **Deleterious** |
| G777I | -0.064 | Neutral | E110M | -2.008 | Neutral |
| **G104H** | **-4.797** | **Deleterious** | **D767M** | **-2.697** | **Deleterious** |
| S105H | -2.381 | Neutral | R768M | -1.790 | Neutral |
| **S106H** | **-3.091** | **Deleterious** | K769M | -1.817 | Neutral |
| **V107H** | **-2.902** | **Deleterious** | K770M | -1.885 | Neutral |
| **L108H** | **-5.866** | **Deleterious** | K771M | -1.363 | Neutral |
| S109H | -2.408 | Neutral | N772M | -0.699 | Neutral |
| E110H | -1.894 | Neutral | K773M | -0.919 | Neutral |
| D767H | -1.866 | Neutral | A774M | -0.765 | Neutral |
| R768H | -1.793 | Neutral | R775M | -0.410 | Neutral |
| K769H | -1.430 | Neutral | S776M | -0.462 | Neutral |
| K770H | -1.945 | Neutral | G777M | 0.196 | Neutral |
| K771H | -1.354 | Neutral | **G104F** | **-6.098** | **Deleterious** |
| N772H | -0.563 | Neutral | **S105F** | **-3.539** | **Deleterious** |
| K773H | -0.885 | Neutral | **S106F** | **-3.327** | **Deleterious** |
| A774H | -0.955 | Neutral | V107F | -2.467 | Neutral |
| R775H | -0.618 | Neutral | **L108F** | **-3.621** | **Deleterious** |
| G777H | 0.311 | Neutral | **S109F** | **-3.937** | **Deleterious** |
| **G104R** | **-4.900** | **Deleterious** | **E110F** | **-3.135** | **Deleterious** |
| S105R | -1.060 | Neutral | **D767F** | **-2.986** | **Deleterious** |
| **S106R** | **-2.574** | **Deleterious** | **R768F** | **-2.687** | **Deleterious** |
| **V107R** | **-2.525** | **Deleterious** | **K769F** | **-2.614** | **Deleterious** |
| **L108R** | **-4.953** | **Deleterious** | **K770F** | **-2.771** | **Deleterious** |
| S109R | -2.281 | Neutral | K771F | -1.673 | Neutral |
| E110R | -1.694 | Neutral | N772F | -1.214 | Neutral |
| D767R | -2.320 | Neutral | K773F | -1.447 | Neutral |
| K769R | -0.360 | Neutral | A774F | -1.122 | Neutral |
| K770R | -0.669 | Neutral | R775F | -0.978 | Neutral |
| K771R | -0.638 | Neutral | S776F | -0.740 | Neutral |
| N772R | -0.410 | Neutral | G777F | -0.095 | Neutral |
| K773R | -0.398 | Neutral | **G104W** | **-5.638** | **Deleterious** |
| A774R | -0.712 | Neutral | **S105W** | **-4.037** | **Deleterious** |
| G777R | 0.219 | Neutral | **S106W** | **-4.204** | **Deleterious** |
| **G104Q** | **-4.358** | **Deleterious** | **V107W** | **-3.864** | **Deleterious** |
| S105Q | -1.503 | Neutral | **L108W** | **-5.385** | **Deleterious** |
| S106Q | -2.210 | Neutral | **S109W** | **-4.463** | **Deleterious** |
| V107Q | -2.206 | Neutral | **E110W** | **-3.686** | **Deleterious** |
| **L108Q** | **-4.819** | **Deleterious** | **D767W** | **-2.935** | **Deleterious** |
| S109Q | -1.460 | Neutral | **R768W** | **-2.822** | **Deleterious** |
| E110Q | -0.750 | Neutral | K769W | -2.391 | Neutral |
| D767Q | -1.632 | Neutral | **K770W** | **-2.696** | **Deleterious** |
| K769Q | -1.029 | Neutral | K771W | -1.887 | Neutral |
| K770Q | -1.194 | Neutral | N772W | -1.288 | Neutral |
| K771Q | -1.026 | Neutral | K773W | -1.393 | Neutral |
| N772Q | -0.199 | Neutral | A774W | -1.115 | Neutral |
| K773Q | -0.209 | Neutral | R775W | -1.060 | Neutral |
| A774Q | -0.440 | Neutral | S776W | -0.975 | Neutral |
| R775Q | -0.002 | Neutral | G777W | -0.368 | Neutral |
| G777Q | 0.499 | Neutral | **G104A** | **-3.809** | **Deleterious** |
| **G104D** | **-2.955** | **Deleterious** | S105A | -1.180 | Neutral |
| S105D | -2.124 | Neutral | S106A | 0.838 | Neutral |
| **S106D** | **-2.559** | **Deleterious** | V107A | 0.739 | Neutral |
| **V107D** | **-2.833** | **Deleterious** | **L108A** | **-4.238** | **Deleterious** |
| **L108D** | **-6.508** | **Deleterious** | S109A | -1.448 | Neutral |
| S109D | -0.953 | Neutral | E110A | -0.903 | Neutral |
| E110D | -0.659 | Neutral | D767A | -2.293 | Neutral |
| D767D | 0.000 | Neutral | R768A | -1.702 | Neutral |
| K769D | -1.751 | Neutral | K769A | -1.677 | Neutral |
| K770D | -1.919 | Neutral | K770A | -1.959 | Neutral |
| K771D | -1.430 | Neutral | K771A | -1.569 | Neutral |
| N772D | -0.461 | Neutral | N772A | -0.733 | Neutral |
| K773D | -0.677 | Neutral | K773A | -1.003 | Neutral |
| A774D | -0.533 | Neutral | A774A | 0.000 | Neutral |
| R775D | -0.624 | Neutral | R775A | -0.345 | Neutral |
| G777D | 0.790 | Neutral | S776A | -0.307 | Neutral |
| **G104N** | **-3.275** | **Deleterious** | G777A | 0.386 | Neutral |
| S105N | -1.381 | Neutral | **G104C** | **-6.287** | **Deleterious** |
| S106N | -2.369 | Neutral | **S105C** | **-2.958** | **Deleterious** |
| V107N | -2.385 | Neutral | S106C | -2.063 | Neutral |
| **L108N** | **-5.767** | **Deleterious** | V107C | -1.597 | Neutral |
| S109N | -1.249 | Neutral | **L108C** | **-4.505** | **Deleterious** |
| E110N | -1.184 | Neutral | **S109C** | **-3.216** | **Deleterious** |

**Table 2.** Prediction of whether the substitution or deletion of an amino acid affects the protein's biological function (ACE-2:NP\_068576.1) by PROVEAN tool.
|  |  |  |  |  |  |  |
| --- | --- | --- | --- | --- | --- | --- |
|  | D767N | -1.180 | Neutral | **E110C** | **-2.939** | **Deleterious** |
|  | R768N | -1.654 | Neutral | **D767C** | **-2.974** | **Deleterious** |
|  | K769N | -1.337 | Neutral | **R768C** | **-2.593** | **Deleterious** |
|  | K770N | -1.576 | Neutral | **K769C** | **-2.584** | **Deleterious** |
|  | K771N | -0.935 | Neutral | **K770C** | **-2.729** | **Deleterious** |
|  | N772N | 0.000 | Neutral | K771C | -2.101 | Neutral |
|  | K773N | -0.551 | Neutral | N772C | -1.364 | Neutral |
|  | A774N | -0.759 | Neutral | K773C | -1.536 | Neutral |
|  | R775N | -0.460 | Neutral | A774C | -0.845 | Neutral |
|  | S776N | -0.189 | Neutral | R775C | -0.795 | Neutral |
|  | G777N | 0.829 | Neutral | S776C | -0.947 | Neutral |
|  | **G104del** | **-9.872** | **Deleterious** | G777C | -0.249 | Neutral |
|  | **S105del** | **-8.259** | **Deleterious** | **G104Y** | **-5.964** | **Deleterious** |
|  | **S106del** | **-8.259** | **Deleterious** | **S105Y** | **-3.177** | **Deleterious** |
|  | **V107del** | **-7.841** | **Deleterious** | **S106Y** | **-3.356** | **Deleterious** |
|  | **L108del** | **-9.391** | **Deleterious** | V107Y | -2.491 | Neutral |
|  | **S109del** | **-8.394** | **Deleterious** | L108Y | **-4.436** | **Deleterious** |
|  | **E110del** | **-8.128** | **Deleterious** | **S109Y** | **-3.516** | **Deleterious** |
|  | **D767del** | **-4.631** | **Deleterious** | **E110Y** | **-2.727** | **Deleterious** |
|  | **R768del** | **-3.193** | **Deleterious** | **D767Y** | **-2.763** | **Deleterious** |
|  | K769del | -2.450 | Neutral | R768Y | -2.280 | Neutral |
|  | K770del | -2.450 | Neutral | K769Y | -2.172 | Neutral |
|  | K771del | -2.450 | Neutral | K770Y | -2.379 | Neutral |
|  | N772del | -2.018 | Neutral | K771Y | -1.409 | Neutral |
|  | K773del | -1.691 | Neutral | N772Y | -1.038 | Neutral |
|  | A774del | -1.436 | Neutral | K773Y | -1.011 | Neutral |
|  | R775del | -1.089 | Neutral | A774Y | -0.928 | Neutral |
|  | S776del | -0.829 | Neutral | R775Y | -0.797 | Neutral |
|  | G777del | -0.343 | Neutral | S776Y | -0.308 | Neutral |
|  |  |  |  | G777Y | 0.093 | Neutral |

Since all results of G104 and L108 points are negative, these results are used in Table 3 and Table 4 for further analysis. Prediction of protein stability changes based on single-site mutation results predicted by I-Mutant2.0(Emidio et al., 2005)is shown in Table 3. According to the results, it is predicted that single-site mutations of Glycine(G) at position 104 amino acid to other amino acids cause both decreases and increase in protein stability. G104I, G104A, G104P, G104M, G104Y, G104I, G104N, G104L, G104E predicted stability increased remaining G104S, G104D, G104R, G104W, G104T, G104F, G104H, G104Q, G104K, G104C stability decreased. Nevertheless, this was not the case for the Leucine at position 108 because protein stability changes were predicted as decreased overall single-site mutations at the L108 position. RI (Reliability Index) of the decreases is relatively high at L108 if compared to G104*X*(X represents all amino acids.).L108S, L108G, L108K have the highest indexes of reliability for decreasing stability of amino acid. On the other hand, G104T, G105 are the most high indexes of reliability for decreasing stability. All other results are shown in Table 3.

**Table 3.** Prediction of protein stability changes upon single-site mutations results predicted by I-Mutant2.0. (RI: Reliability Index) Temperature in Celsius degree was set 25 and pH(-log[H+]) was set 7 on I-Mutant2.0.

| Used tool | Variant | Stability | RI | Variant | Stability | RI |
| --- | --- | --- | --- | --- | --- | --- |
| I-Mutant2.0 | G104S | Decrease | 5 | G104C | Decrease | 0 |
|  | G104A | Increase | 3 | G104P | Increase | 1 |
|  | G104D | Decrease | 1 | G104M | Increase | 1 |
|  | G104R | Decrease | 3 | G104Y | Increase | 1 |
|  | G104W | Decrease | 2 | G104L | Increase | 0 |
|  | G104I | Increase | 4 | G104K | Decrease | 4 |
|  | G104N | Increase | 0 | G104Q | Decrease | 2 |
|  | G104T | Decrease | 5 | G104V | Increase | 2 |
|  | G104F | Decrease | 1 | G104H | Decrease | 5 |
|  | G104E | Increase | 5 | L108A | Decrease | 8 |
|  | L108F | Decrease | 6 | L108C | Decrease | 7 |
|  | L108D | Decrease | 7 | L108M | Decrease | 6 |
|  | L108S | Decrease | 9 | L108P | Decrease | 5 |
|  | L108R | Decrease | 8 | L108W | Decrease | 4 |
|  | L108G | Decrease | 9 | L108Y | Decrease | 4 |
|  | L108I | Decrease | 6 | L108T | Decrease | 8 |
|  | L108V | Decrease | 8 | L108H | Decrease | 8 |
|  | L108K | Decrease | 9 | L108Q | Decrease | 8 |
|  | L108N | Decrease | 7 | L108E | Decrease | 6 |

**Table 4.** Results of wild-type and mutant-type amino acid (ACE-2: NP_068576.1))properties obtained from Project Hope software. ^a^Could not be retrieved.

| Variant | WILD-TYPE AMINO ACIDS |  |  | MUTANT -TYPE AMINO ACIDS |  |  |  |  |
| --- | --- | --- | --- | --- | --- | --- | --- | --- |
|  | Size | Charge | Hydrophobicity | Size | Charge | Hydrophobicity | Mutant Residue | Damaging to the protein conservation |
| G104S | < | a | a | > | a | a | Might lead to bumps | Probably Yes |
| G104A | < | a | < | > | a | > | Might lead to bumps | Probably Yes |
| G104D | < | Neutral | > | > | Negative | < | Might lead to bumps | **Probably No** |
| G104R | < | Neutral | > | > | Positive | < | Might lead to bumps | Probably Yes |
| G104W | < | <sup>a</sup> | < | > | <sup>a</sup> | > | Might lead to bumps | Probably Yes |
| G104I | < | <sup>a</sup> | < | > | <sup>a</sup> | > | Might lead to bumps | Probably Yes |
| G104N | < | <sup>a</sup> | > | > | <sup>a</sup> | < | Might lead to bumps | Probably Yes |
| G104T | < | <sup>a</sup> | <sup>a</sup> | > | <sup>a</sup> | <sup>a</sup> | Might lead to bumps | Probably Yes |
| G104F | < | <sup>a</sup> | < | > | <sup>a</sup> | > | Might lead to bumps | Probably Yes |
| G104E | < | Neutral | > | > | Negative | < | Might lead to bumps | **Probably No** |
| G104C | < | <sup>a</sup> | < | > | <sup>a</sup> | > | Might lead to bumps | Probably Yes |
| G104P | < | <sup>a</sup> | < | > | <sup>a</sup> | > | Might lead to bumps | Probably Yes |
| G104M | < | <sup>a</sup> | < | > | <sup>a</sup> | > | Might lead to bumps | Probably Yes |
| G104Y | < | <sup>a</sup> | <sup>a</sup> | > | <sup>a</sup> | <sup>a</sup> | Might lead to bumps | **Probably No** |
| G104L | < | <sup>a</sup> | < | > | <sup>a</sup> | < | Might lead to bumps | Probably Yes |
| G104K | < | Neutral | > | > | Positive | < | Might lead to bumps | Probably Yes |
| G104Q | < | <sup>a</sup> | > | > | <sup>a</sup> | < | Might lead to bumps | **Probably No** |
| G104V | < | <sup>a</sup> | < | > | <sup>a</sup> | > | Might lead to bumps | Probably Yes |
| G104H | < | <sup>a</sup> | > | > | <sup>a</sup> | < | Might lead to bumps | Probably Yes |
| L108F | < | <sup>a</sup> | <sup>a</sup> | > | <sup>a</sup> | <sup>a</sup> | Might lead to bumps | **Probably No** |
| L108D | < | Neutral | > | > | Positive | < | Might lead to bumps | Probably Yes |
| L108S | > | <sup>a</sup> | > | < |  | < | Might lead to loss of interactions | Probably Yes |
| L108R | < | Neutral | > | > | Positive | < | Might lead to bumps | Probably Yes |
| L108G | > | <sup>a</sup> | > | < | <sup>a</sup> | < | Might lead to bumps | Probably Yes |
| L108I | = | <sup>a</sup> | <sup>a</sup> | = | <sup>a</sup> | <sup>a</sup> | **Not given** | Probably Yes |
| L108V | > | <sup>a</sup> | <sup>a</sup> | < | <sup>a</sup> | <sup>a</sup> | Might lead to loss of interactions | Probably Yes |
| L108K | < | Neutral | > | > | Positive | < | Might lead to bumps | Probably Yes |
| L108N | < |  | > | > | <sup>a</sup> | < | Might lead to bumps | Probably Yes |
| L108A | > | <sup>a</sup> | <sup>a</sup> | < | <sup>a</sup> | <sup>a</sup> | Might lead to loss of interactions | Probably Yes |
| L108C | > | <sup>a</sup> | <sup>a</sup> | < | <sup>a</sup> | <sup>a</sup> | Might lead to loss of interactions | Probably Yes |
| L108M | < | <sup>a</sup> |  | > | <sup>a</sup> | <sup>a</sup> | Might lead to bumps | Probably Yes |
| L108P | > | <sup>a</sup> | <sup>a</sup> | < | <sup>a</sup> | <sup>a</sup> | Might lead to loss of interactions | Probably Yes |
| L108W | < | <sup>a</sup> | <sup>a</sup> | > | <sup>a</sup> | <sup>a</sup> | Might lead to bumps | Probably Yes |
| L108Y | < | <sup>a</sup> | > | > | <sup>a</sup> | < | Might lead to bumps | Probably Yes |
| L108T | > | <sup>a</sup> | > | < | <sup>a</sup> | < | Might lead to loss of interactions | Probably Yes |
| L108H | < | <sup>a</sup> | > | > | <sup>a</sup> | < | Might lead to bumps | Probably Yes |
| L108Q | < | <sup>a</sup> | > | > | <sup>a</sup> | < | Might lead to bumps | Probably Yes |
| L108E | < | Neutral | > | > | Negative | < | Might lead to bumps | **Probably No** |

Each amino acid has its own specific size, charge, and hydrophobicity-value. The original wild-type residue and newly introduced mutant residue often differ in these properties(Hanka et al.,2010). The results of Table 4 are obtained based on a report that will evaluate the effect of the mutation on the following features: Contacts made by the mutated residue, structural domains in which the residue is located, modifications on this residue, and known variants for this residue(Table 4). A feature will only be shown when information is available. A short conclusion based on just the amino acid properties is always shown(Hanka et al.,2010). If the mutants residues are bigger, this might lead to bumps. Except L108S,L108G,L108I,L108V,L108A,L108C, all of other variant residue are bigger than wild type residue(Table 4). Both mutant residue and wild-type residue are equal to each other for L108I, so the result of this equality is not found (Table 4). For L108S,L108G, L108V,L108A,L108C, the mutants residues are smaller, might lead to loss of interactions for ACE-2 protein (NCBI Acc.No: NP_068576.1)(Table 4). Features of Hydrophobicity are found too. Results are found that their mutant residue can be more or less hydrophobic than the wild-type residue or not found any information about them(Table 4). G104R, G104K, L108D, L108R, L108K wild types residue charge were Neutral, and the mutant types residue charge were Positive while G104D, G104E, L108E wild types residue charge were Neutral, the mutant types residue charge was Negative(Table 4). Except for these regions, other charges are not predicted by Project Hope software.

When the wild-type residue is glycine, the most flexible of all residues, this flexibility might be necessary for the protein’s function. This feature was not found for the L108 position. Mutation of this glycine can abolish this function based on the Hope tool’s result(Hanka et al.,2010). Except for G104D, G104E, G104Y, G104Q, L108F, L108E all of the other variant which is belongs to G104 does not damage to ACE-2 protein conservation(Table 4). The finding reason why G104D, G104E, G104Y, G104Q are not damaging to ACE-2 is that the mutant residue is among the other residue types observed at this position in homologous sequences. L108F, L108E does not damage ACE-2 protein conservation because homologous proteins exist with the same residue type as our mutant. Both mean that this mutation can occur at this position is probably not damaging to the protein(Hanka et al.,2010).

The number of amino acids is constant to 805 for all proteins except two proteins which are G108del, L108del. ACE-2 protein (NP_068576.1) molecular weight was found 92463.04kDa. The largest one was G104W with~92592 kDa, whereas L108del with~92350 kDa was the smallest (Table 5). When all variants include normal protein, variants, and deletion types encoded by the full genome, were analyzed, the theoretical PI value was between 5.32 and 5.41. The theoretical PI of the normal one was 5.36. Among normal and variant proteins, all were negatively charged(Table 5). The estimated half-lives were 30 hours for all proteins in Mammalian reticulocyte (in vitro),> 20 hours in yeast(in vivo), >10 hours in E.coli(in vivo). All variants instability index were found as instable except L108G(39.98/stable), L108Y(40.0/stable), L108T(39.98/stable). Thereof these stable variants are highlighted in black in Table 5. The greatest value is 40.82/unstable, while the smallest one is 39.98/stable. The aliphatic index showed a significant variation ranging between 80.06 to 81.03 among all variants proteins, while the normal one’s aliphatic index was found at 80.55. Nevertheless, overall weighted results of the aliphatic index 80.55 and 80.06 were found(Table 5). The grand average of hydropathicity(GRAVY) value was found negative in all variants, and the normal one’s value was found −0.375. The GRAVY values were between −0.369 and - 0.384(Table 5).

**Tablo 5.**
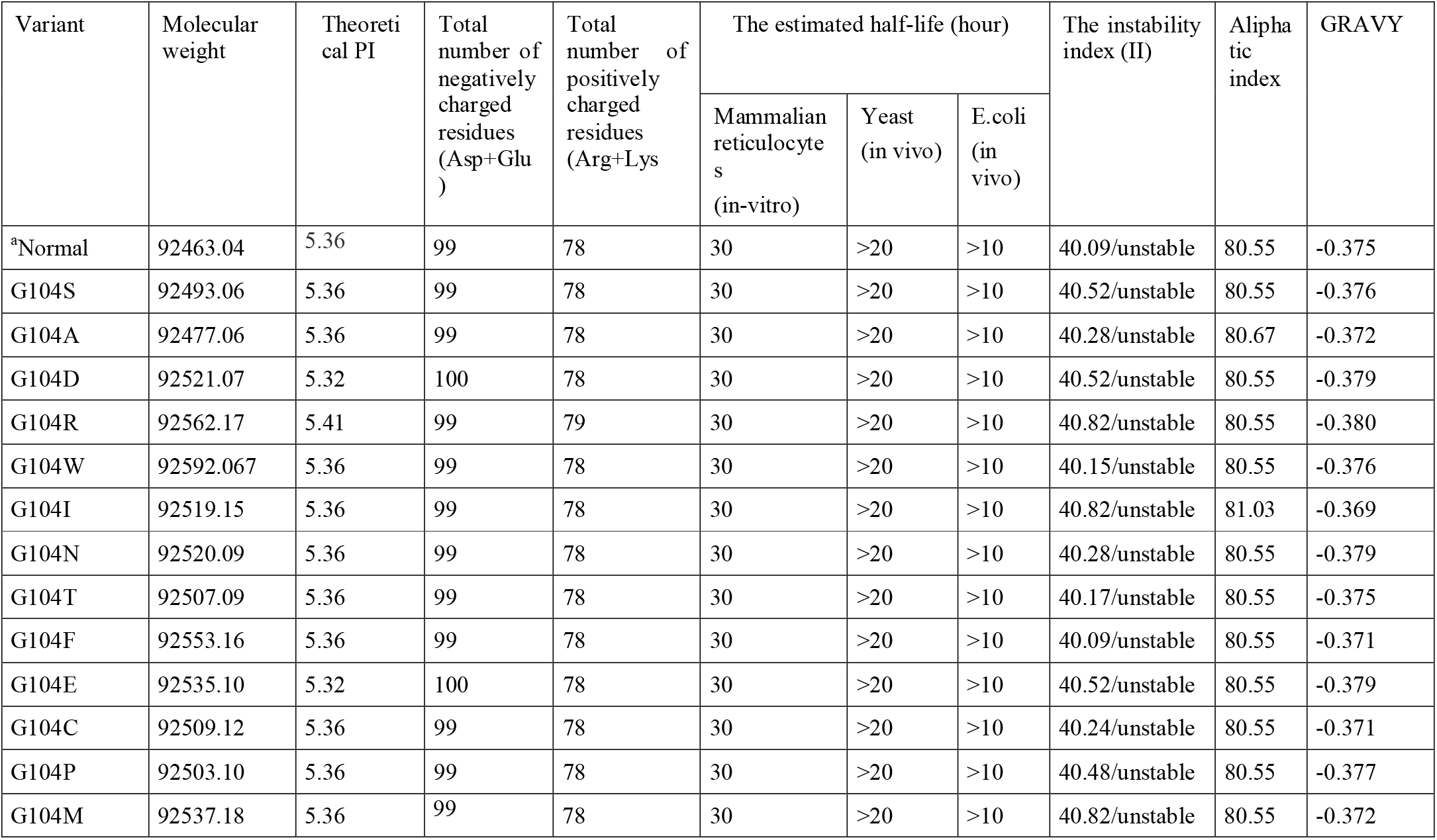
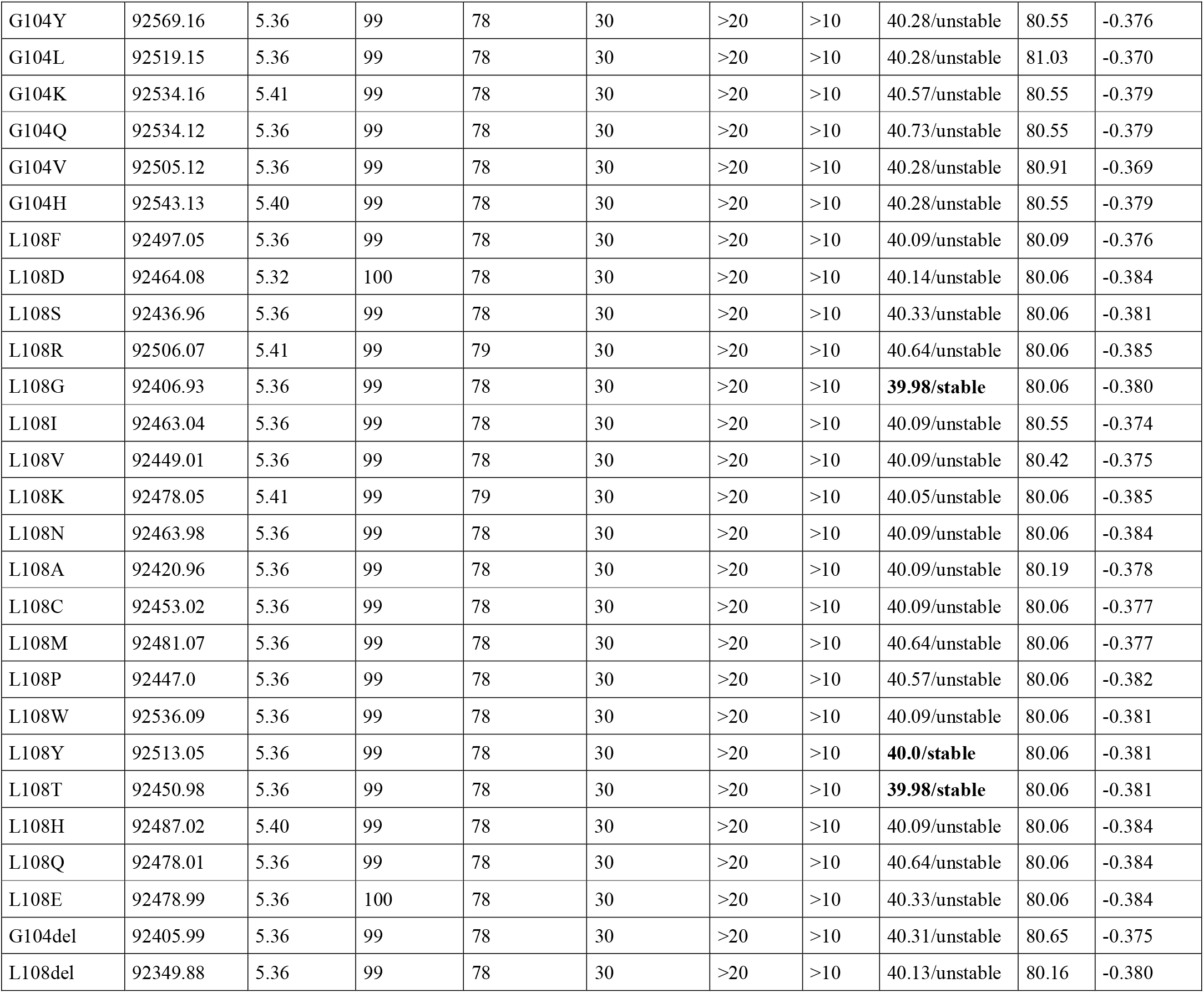
Physico-chemical parameter results predicted by ExPASyProtParam. ^a^Normal is NP_068576.1

Regions G104 and L108 do not belong to any conserved amino acid of ACE-2 protein based on NP_068576.1 from NCBI (Figure 1). Amino Acids (located on ACE-2 receptor samples) conservation scores results are G(score:1),S(score:9),L(score:3), V(score:8),N(score:7),Q(score:7) while the maximum score is 9 based on ConSurf Server(Ashkenazy et al., 2016). Moreover, G104 (score: 1)is the most susceptible amino acid, and the amino acid L108(score between 3-4) is located far from the conserved site(Figure 1). Variable, unknown, and conserved amino acids are shown for both back(B) and front(A) of ACE-2

**Figure 1.**
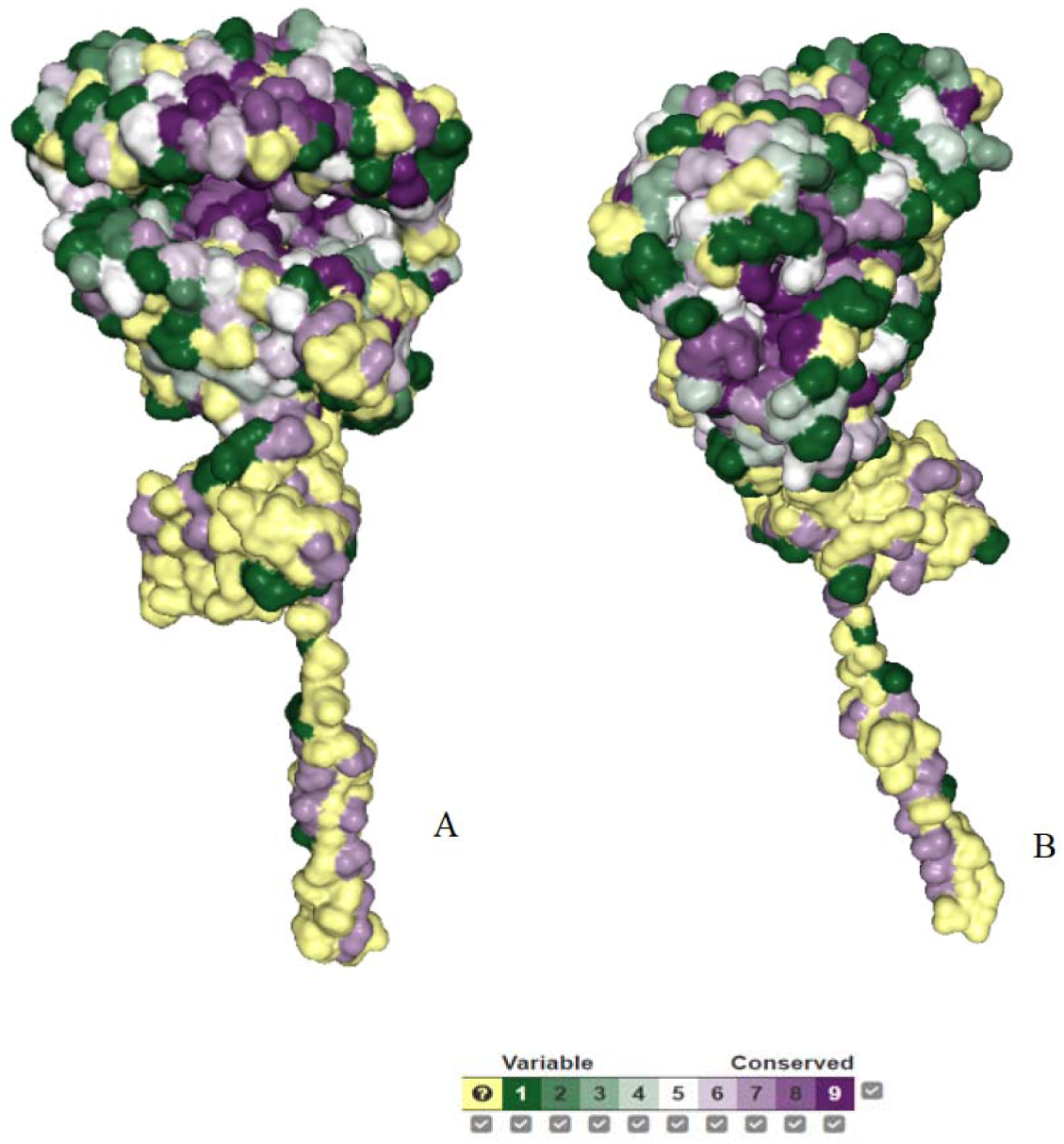
Screenshot of ACE-2’amino acid conservation score with amino acid location results predicted by ConSurf Server. **(A)** is a screenshot of ACE-2’s front while **(B)** shows the screenshot of ACE-2’back site. Variable, unknown, and conserved amino acids are shown for both the back and front of ACE-2 protein. Amino acids (located between 101 and 109 on ACE-2 receptor samples) conservation scores results are G(1), S(9), L(3), V(8), N(7), Q(7), while the maximum results are 9.

The tertiary structure of LQQNGSSVL(A) was predicted by 3Dpro(Gaëlle et al.,2009), and its visualization was obtained by Chimera 1.15(Figure 2)((Pettersen et al.,2004).). Docking of LQQNGSSVL with both ACE-2 protein (B) and SARS CoV’s surface protein(S) are shown(C) (Figure 2).

**Figure 2.**
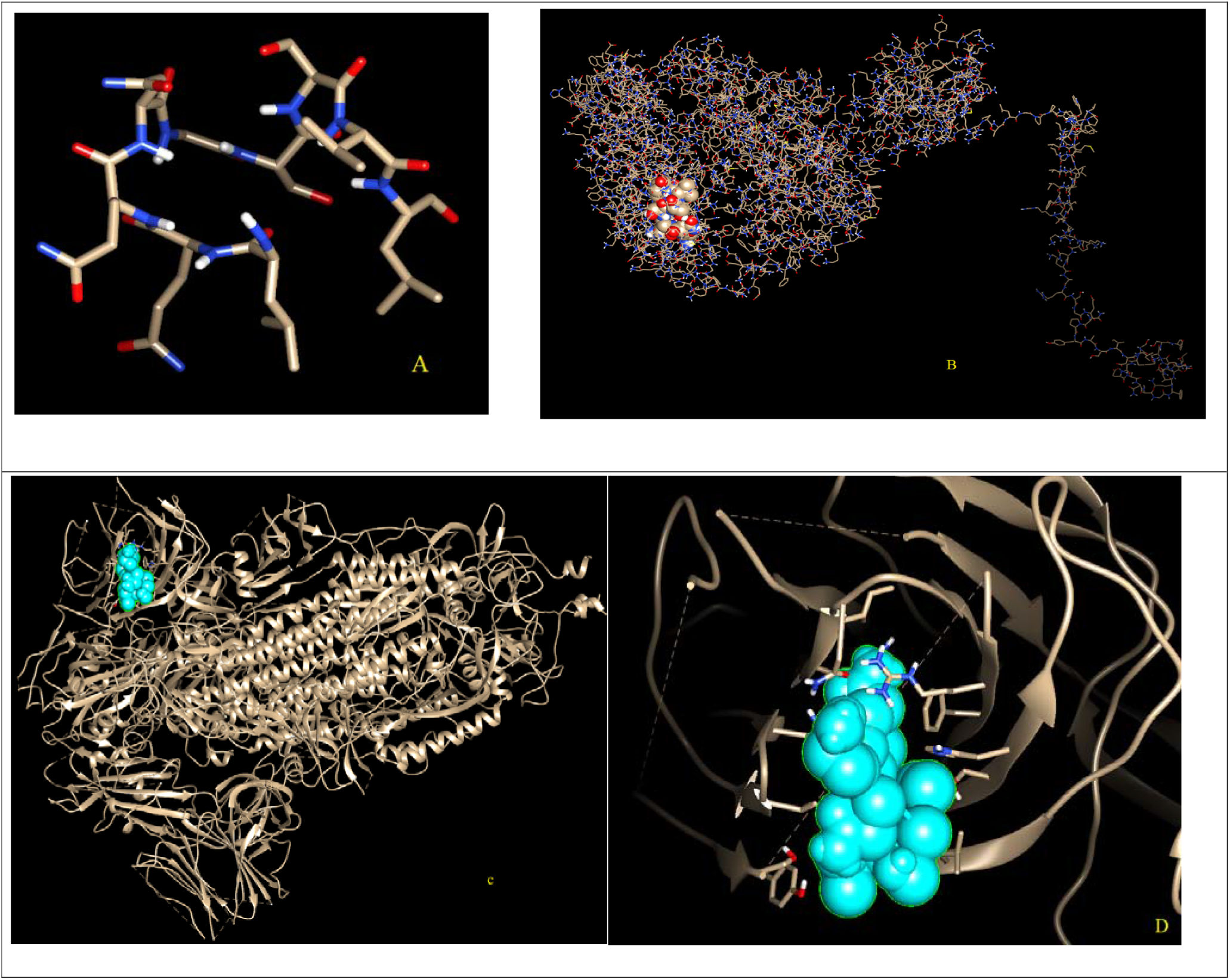
**(A)** Tertiary structure of LQQNGSSVL (epitope) as PDB predicted by 3Dpro.**(B)** the snapshot representing our interested sequence location in the pocket of the molecular surface of the ACE-2 protein. **(C)** Predicted where LQQNGSSVL (Cyan)epitope and DSSVLSEDK(Cyan) epitope bind to SARS CoV-2 surface protein(pdb:6VXX). **(D)** docking between Spike (pdb:6VXX) and LQQNGSSVL (all the structures are visualized using Chimera 1.15).

The predicted interaction of wild-type ACE-2(NP_068576.1) with spike protein(6VXX) is shown in Figure 3(A). While the yellow color has the best-predicted docking score (−261.75), the green color has the other best-predicted docking score(−247.04) in the 10 top models for wilt-type ACE-2(Table 6). The predicted interaction mutant ACE-2(G104del) with surface glycoprotein (6VXX) is shown in Figure 3(B). The best-predicted docking score(The yellow one) for this interaction is −241.10, and the other best-predicted docking score(green) is - 240.87 in 10 top models(Table 6). The predicted interaction in the form of ACE-2(L108del) with spike protein(pdb:6VXX) result is in Table 6. The best-predicted docking score(the yellow one) for ACE-2(L108del) is −248.11, and the other best-predicted docking score(green) is - 235.25 in the 10 predicted top models**(Fig.3-C)**. The predicted interaction between a form of ACE-2(coffee), which has both G104del and L108del, and spike protein result is shown in **Figure 3(D)**. The yellow one shows that the best docking score(−257.44), the green shows that the other best docking score(−254.48) is predicted in 10 top models**(Figure 3(D))**.

**Figure 3.**
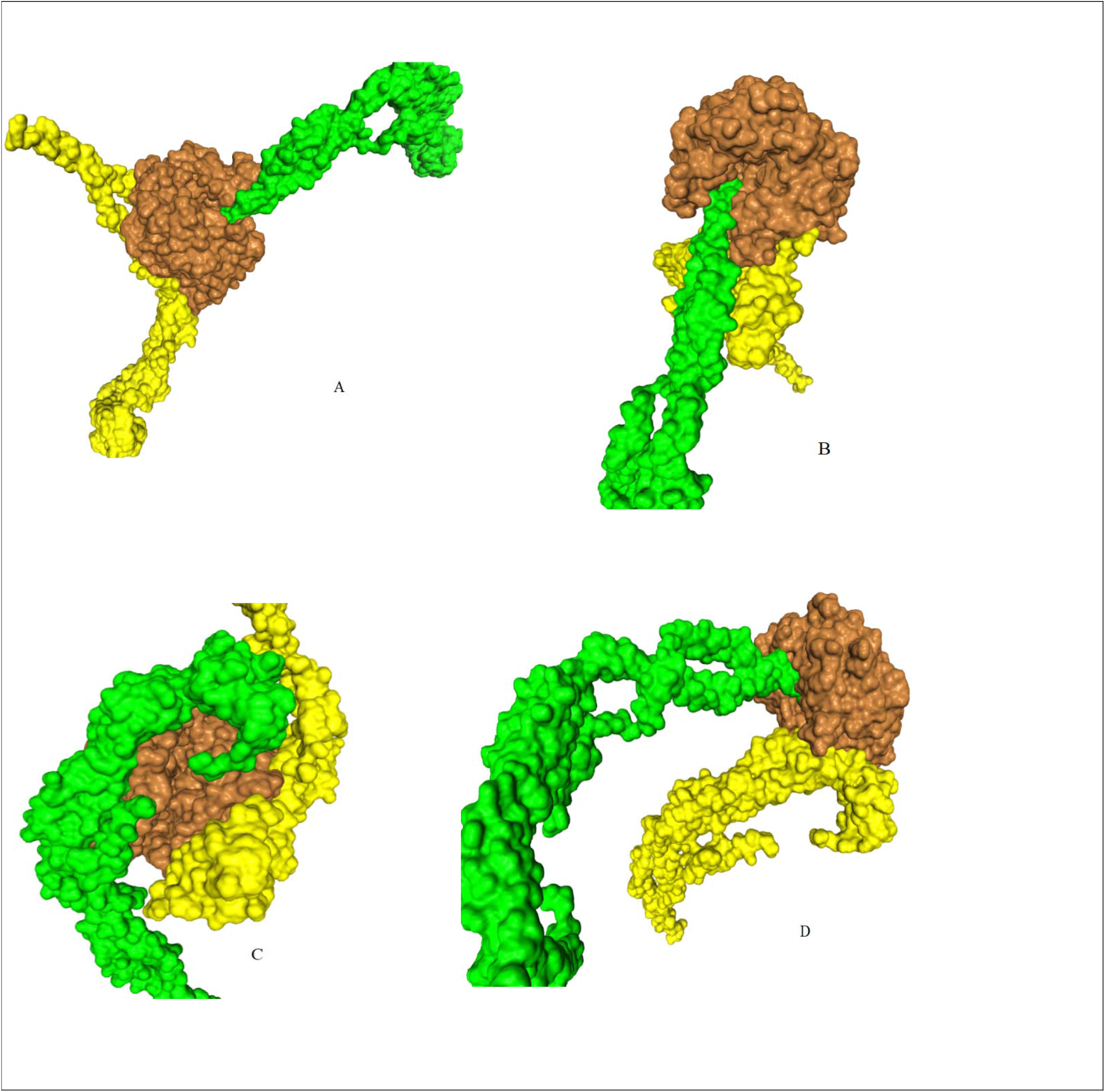
Screenshots of two best models of in the 10 top predicted models of interaction between spike protein(pdb:6VXX) -ACE-2 receptor(NP_068576.1). The normal ACE-2 **(A),** G104del**(B),** L108del**(C),** both G104del and L108del**(D)** results were predicted by HDOCK SERVER.( Brown represents ACE-2 receptor, while yellow (has the best score) and green(has the other best docking score) represent the best two alternatives to the predicted sars protein)

**Table 6.** The two highest predicted docking scores (between ACE-2 receptor and spike protein)of in the top ten models results obtained by HDOCK SERVER. The least docking scores are highlighted in black

| Variant | Two predicted docking scores of the top ten models |  | Variant | Two predicted docking scores of the top ten models |  |
| --- | --- | --- | --- | --- | --- |
|  | First model | Second model |  | First model | Second model |
| Normal | -261.75 | -247.04 | Normal | -261.75 | -247.04 |
| G104S | -258.21 | -253.13 | L108F | -279.14 | -248.37 |
| G104A | **-234.30** | **-231.27** | L108D | -252.92 | -245.59 |
| G104D | -246.23 | -245.81 | L108S | -253.92 | -253.01 |
| G104R | -249.56 | -231.95 | L108R | -274.68 | -242.54 |
| G104W | -250.24 | -239.80 | L108G | -269.34 | -237.52 |
| G104I | -247.65 | -246.44 | L108I | -241.86 | -239.13 |
| G104N | -251.46 | -245.32 | L108V | -263.55 | -245.52 |
| G104T | -252.48 | -243.87 | L108K | -246.54 | -245.02 |
| G104F | -250.53 | -241.69 | L108N | -261.75 | -247.04 |
| G104E | -247.59 | -243.56 | L108A | -244.93 | -234.17 |
| G104C | -248.39 | -245.50 | L108C | -256.33 | -229.17 |
| G104P | -249.40 | -247.01 | L108M | -252.04 | -251.34 |
| G104M | -258.29 | -244.11 | L108P | -255.39 | -240.77 |
| G104Y | -258.41 | -250.82 | L108W | -260.16 | -246.76 |
| G104L | **-234.64** | **-232.41** | L108Y | -287.06 | -227.04 |
| G104K | -256.61 | -244.67 | L108T | -249.22 | -247.40 |
| G104Q | -266.53 | -240.85 | L108H | -259.13 | -251.41 |
| G104V | -261.75 | -247.04 | L108Q | -246.83 | -243.59 |
| G104H | -261.75 | -247.04 | L108E | -245.08 | -239.35 |
| G104del | -241.10 | -240.87 | L108del | -248.11 | - 235.25 |

## Discussion

This study aimed to make a detailed bioinformatics analysis of the ACE-2 receptor(NP_068576.1). Examination of ACE-2’structure may play a vital role in understanding how it works(Li et al.,2008; Ankita et al.,2017;Can et al.,2020; Aktas el al.,2021). Therefore, unregulated/unstructured regions of ACE-2 protein results predicted by the disEMBL tool(Table 1). This tool has three different definitions, which are disordered by Loops/coils definition, disordered by Hot-loops definition, and disordered by Remark-465 definition(Linding et al.,2003). The regions of three different definitions of the tools slightly differ as they both work on a different algorithm and principle(Linding et al.,2003;Ankita et al.,2017). After these three definitions, the common regions’ amino acids shown in the upper case letters indicate the disordered regions; therefore, they are chosen(Table 1). All these results have common values that have encouraged us to work on places selected from these regions. Based on sequence of ACE-2(NP_068576.1), the predicted unregulated/unstructured regions are GSSVLSE(104-110) and DRKKKNKARS(767-777) respectively(Table 1).

These GSSVLSE(104-110) and DRKKKNKARS(767-777) regions were analyzed with PROVEAN(Protein Variation Effect Analyzer) tool to predicted for more detailed analysis and research effect on the biological function of the protein(Choi et al.,2012). All amino acids (NP_068576.1) in this GSSVLSE (104-110) and DRKKKNKARS (767-777) region were point mutated one by one. The aim is that predict whether the substitution or deletion of an amino acid effects on the biological function of the protein (NP_068576.1)(Table 2)( Choi et al.,2012). According to the PROVEAN tool’s result, it was determined that all the results obtained at the G104 and L108 points negatively affect the biological activity of ACE-2 protein. Furthermore, it also was predicted that both G104 and L108 do not belong to conservation regions of ACE-2 protein(Ashkenazy et al.,2016). The results of both point mutations and deletions were found to be deleterious with remarkable results which variants with the result of the PROVEAN tool score less than or equal to −2.5 are considered “deleterious” (Table 2) ( Choi et al.,2012; Yongwook et al., 2015). While the most harmless result was G777del(score:-0.343), the most harmful was predicted as G104del(score:-9.872)(Table 2). There are regions other than G104 and L108 with negative results(Table 2), but we wanted to continue on these two points. Because in the case of all possible mutations and deletions at these points, the results are deleterious(Table 2), leading to the idea that this place may play a critical role for the ACE-2 receptor. Besides, it was stated in a study that a protein is effective in signaling pathways of biological function(Linden,2017). Changes in biological functions can have negative consequences. Therefore, the results we found can be considered to be very important(Hong et al.,2016; Linden,2017).

The I-Mutant2.0 tool was used to estimate protein stability changes based on single point mutation results in both the G104 and L108 regions. The results here made it even more meaningful for us to continue in these two regions (Table 3). According to the results, it is predicted that single-site mutations of Glycine(G) at position 104 amino acid to other amino acids cause both decreases and increase in protein stability. G104I, G104A, G104P, G104M, G104Y, G104I, G104N, G104L, G104E predicted stability increased remaining G104S, G104D, G104R, G104W, G104T, G104F, G104H, G104Q, G104K, G104C stability decreased. However, this was not the case for the Leucine at position 108 because protein stability changes were predicted as decreased overall single-site mutations at the L108 position(Table 3). In this case, it can be considered that the L108 is more effective than G104 in the stability of the protein of interest. Because it was seen that all possible mutations and deletions in the L108 region result in stability decreased. It is known that stability is a fundamental property affecting the function, activity, and regulation of biomolecules. Stability changes are often found for mutated proteins involved in diseases (Sofia et al.,2010). Based on this study(Sofia et al.,2010), therefore, the results in Table 3 may trigger some diseases.

It is stated that charge and hydrophobicity play a vital role in the toxicity of protein and macroscopic properties(Kelsey et al.,2013; Mannini et al.,2014). Each amino acid has its own specific size, charge, and hydrophobicity-value(Hanka et al.,2010). It was found that possible mutational changes in the G104 and L108 regions may cause changes in the charge, size, and hydrophobicity of amino acids(Table 4). Based on the study(Kelsey et al.,2013) done, these changes may be considered essential for protein toxicity. Furthermore, it was determined with the Hope tool that the change in the G104 region affected the flexibility of the ACE-2 protein(Hanka et al.,2010). When the wild-type residue is a glycine, this is the most flexible of all residues. This flexibility might be necessary for the protein’s function, but this feature was not found for the L108 position(Table 4). Therefore, mutation of this glycine can abolish this function based on the result of the Hope tool. Furthermore, when the mutant size is bigger than the wild type, this process might lead to bumps. On the other hand, when the wild type of amino acid is bigger than the mutant, this process was predicted to might lead to loss of interactions(Table 4) (Hanka et al.,2010). Finally, it was determined that some of their mutations damage the conservation of proteins. For these reasons, both the G104 and L108 regions can be considered to be very important for ACE-2 protein(Sofia et al.,2010; Hanka et al.,2010; Kelsey et al.,2013; Mannini et al.,2014; Hong et al.,2016.).

The physical-chemical analysis showed that all proteins, ACE-2, its variant, and deletions(Table 5), had a negative GRAVY value indicating that all proteins are hydrophilic and have a good interaction with the surrounding water molecules(Droppa-Almeida et al.,2018). Also, it had stable, which are important parameters for biophysical studies on epitope-based vaccine design, but only L108G(39.98/stable), L108Y(40.0/stable), L108T(39.98/stable) predicted as stable(Table 5). Moreover, all proteins had a central aliphatic index that shows stability in a broad spectrum of temperature(Shey et al.,2019), fewer than two transmembrane helices easing cloning and expression(Meunier et al.,2016; Can et al.,2020) a considerable molecular weight and long estimated half-life of more than 10 hours for all organisms. The estimated half-life was 30 hours for all proteins in Mammalian reticulocyte (in vitro),> 20 hours in yeast(in vivo), >10 hours in E.coli(in vivo)(Table 5). Another crucial point for these proteins was predicting one posttranslational modification (N glycosylation region)(Table 5). The presence of these modifications may be important for these proteins to obtain by recombinant technology, eukaryotic expression systems, and bacteria(Hansson et al.,2000;Can et al.,2020).

All residues in ACE-2 protein are not equally important. Some are essential for the protein’s proper structure and function, whereas others can be readily replaced(Capra et al.,2007). The evolutionary conservation of an amino acid in the ACE-2 protein is essential because the degree of evolutionary conservation of an amino acid in a protein reflects a balance between its natural bias to mutate and the overall need to retain the structural integrity and function of the protein(Ashkenazy et al.,2016). Based on this study(Ashkenazy et al.,2016), it may be important to know that G104 and L108 do not belong to the conservation of ACE-2 protein(Figure 2) because this helps to understand the natural tendency to mutate and the overall need to retain the structural integrity and function of the ACE-2 protein(Ashkenazy et al.,2016). Many of the amino acids that make up ACE-2 are not known to be conserved or variable regions. Amino acids whose evolutionary processes are unknown are shown in yellow and cover a great deal of space(Figure 2). Therefore, the conservation scores of the amino acid(belongs to ACE-2) may be so important to understand ACE-2 receptor structure(Can et al.,2020;,Ashkenazy et al.,2016) in terms of both interaction between Spike (Figure 1) and other results(Table 1-6). Besides, it is clearly known that the angiotensin-converting enzyme, expressed on the surface of several pulmonary and extra-pulmonary cell types, including cardiac, renal, intestinal, and endothelial cells(Albini et al.,2020). Knowing the conservation score of amino acids creating ACE-2 protein may help these all things(Lensink et al.,2019;Albini et al.,2020). Even the maximum conservation score is 9 (The proportions are given in Figure 2) based on ConSurf Server(Ashkenazy et al.,2016), G104 predicted conservation score is 1, and L108 predicted conservation score of L108 is 3 (Figure 2). The predicted and obtained all results(all tables and figures) about ACE-2 may keep their validity because of not being a conservation region.

The HDOCK server has been compared with many other protein-protein interaction tools, and the results are satisfactory(Yan et al.,2020). Also, this tool has been used in some studies in protein-protein interaction(Amar et al.,2018; Benjamin et al.,2019; Lensink, M. F. et al.,2019). One logic of this tool, all the predicted binding modes are ranked according to their binding energy scores, in which the top-scored models are chosen as the predicted complex structures(Sheng,2014). Therefore, although at least 50 results were found, based on this logic(Sheng,2014), the two highest predicted scores were used(Table 6) as it was predicted that these regions were quite important for ACE-2 protein(Table 1-5). It was determined that the change of amino acids in the G104 and L108 regions affected the ACE-2 protein-spike protein interaction, too(Tablo 6). Based on natural changes(Table 6), changes in G104 and L108 points can produce both positive and negative results(Table 6). According to the results in Table 6, it can be thought that the changes in the G104 region generally decrease the interaction of ACE-2 with the spike protein, while the changes in the L108 region are mostly increasing. The two lowest predicted docking scores at the G104 point belong to the G104A and G104L changes, while the highest predicted docking scores in the L108 region belong to the changes of L108Y and L108F. It was determined that these regions’ changes could also affect the predicted conformation of the ACE2 protein-spike protein interaction(Figure 3). Therefore, these results also support that the amino acids at the points G104 and L108 can play a very active role in the structure, stability, interaction with other proteins, and possible mutations in ACE-2(Table 1-6, Figure 1-3). The results(Table1-6, Figure1-3) of this study are aimed to have positive contributions to both ACE-2-related diseases and ACE-2 _SARS CoV interaction.

## Conclusion

As a result of bioinformatics studies, some critical chemical and physical properties of the ACE-2 receptor were determined. It was determined that especially the G104 and L108 regions are essential for ACE-2 protein. All possible mutations and deletions in these regions affect both the structure of ACE-2 protein and the biological activity of ACE-2 protein. And also, they may play a vital role in interacting with other proteins. It may be thought that ACE-2’amino acid at G104 and L108(both do not belong to the conservation region) plays a crucial role in ACE-2 protein. These results are expected to shed light on both SARS-CoV and ACE-2 protein’s related diseases.

## References

1. Wu, J. T., Leung, K. & Leung, G. M. Nowcasting and forecasting the potential domestic and international spread of the 2019-nCoV outbreak originating in Wuhan, China: a modelling study. Lancet 2020, 395, 689–697 https://doi.org/10.1016/S0140-6736(20)30260-9

2. Chan-Yeung M & Yu WC Outbreak of severe acute respiratory syndrome in Hong Kong Special Administrative Region: a case report. British Medical Journal 2003, 32: 850–852 https://doi.org/10.1136/bmj.326.7394.850

3. Belouzard, S., Chu, V. C and Whittaker, G. R. Activation of the SARS coronavirus spike protein via sequential proteolytic cleavage at two distinct sites. Proc. Natl. Acad. Sci. USA 2009. 5871–5876 https://doi.org/10.1073/pnas.0809524106

4. Hui, D. S. et al. The continuing 2019-nCoV epidemic threat of novel coronaviruses to global health - the latest 2019 novel coronavirus outbreak in Wuhan, China. Intl. J. Infect. Dis. 2020, 91, 264–266 https://doi.org/10.1016/j.ijid.2020.01.009

5. Aktas E. Korcan E., Ozdemir Ozgenturk N, Erismis UC. Bioinformatic analysis reveals that some bacteria may aid SARS-COV-2 spread and entry into host cells (Accepted by Sigma Journal of Engineering and Natural Sciences, In press, 2021)

6. Aktas E. Bioinformatics Analysis Unveils Certain Mutations Implicated in Spike Structure Damage and Ligand-Binding Site of Severe Acute Respiratory Syndrome Coronavirus 2. Bioinformatics and Biology Insights. 2021. https://doi.org/10.1177/11779322211018200

7. Xiaocong Pang, Yimin Cui & Yizhun Zhu Recombinant human ACE2: potential therapeutics of SARS-CoV-2 infection and its complication, Acta Pharmacologica Sinica 2020, volume 41, pages 1255–1257 https://doi.org/10.1038/s41401-020-0430-6

8. Hoffmann M, Kleine-Weber H, Schroeder S, Krüger N, Herrler T, Erichsen S, et al. SARS-CoV-2 cell entry depends on ACE2 and TMPRSS2 and is blocked by a clinically proven protease inhibitör Cell. 2020, Volume 181, Issue 2, Pages 271–280.e8 https://doi.org/10.1016/j.cell.2020.02.052

9. Zou Z, Yan Y, Shu Y, Gao R, Sun Y, Li X, et al. Angiotensin-converting enzyme 2 protects from lethal avian influenza A H5N1 infections. Nat Commun; 2014, 5:3594. https://doi.org/10.1038/ncomms4594

10. Yang P, Gu H, Zhao Z, Wang W, Cao B, Lai C, et al. Angiotensin-converting enzyme 2 (ACE2) mediates influenza H7N9 virus-induced acute lung injury. Sci Rep; 2014, 4:7027. https://doi.org/10.1038/srep07027

11. Qaradakhi T, Gadanec LK, McSweeney KR, Tacey A, Apostolopoulos V, Levinger I, Rimarova K, Egom EE, Rodrigo L, Kruzliak P, Kubatka P, Zulli A. The potential actions of angiotensin-converting enzyme II (ACE2) activator diminazene aceturate (DIZE) in various diseases. Clin Exp Pharmacol Physiol. 2020 May;47(5):751–758. doi: 10.1111/1440-1681.13251. Epub 2020 Jan 28. PMID: 31901211.

12. Li X, Molina-Molina M, Abdul-Hafez A, Uhal V, Xaubet A, Uhal BD. Angiotensin converting enzyme-2 is protective but downregulated in human and experimental lung fibrosis. Am J Physiol Lung Cell Mol Physiol. 2008, 295:L178–85 DOI: 10.1152/ajplung.00009.2008

13. Zhang H, Baker A. Recombinant human ACE2: acing out angiotensin II in ARDS therapy. Crit Care.; 2017, 21:305. https://doi.org/10.1186/s13054-017-1882-z

14. Can, H., Köseoğlu, A.E., Erkunt Alak, S. et al. In silico discovery of antigenic proteins and epitopes of SARS-CoV-2 for the development of a vaccine or a diagnostic approach for COVID-19. Sci Rep 2020, 10, 22387 https://doi.org/10.1038/s41598-020-79645-9.

15. Rashid MI, Rehman S, Ali A, Andleeb S. 2019. Fishing for vaccines against Vibrio cholerae using in silico pan-proteomic reverse vaccinology approach. PeerJ 7:e6223 https://doi.org/10.7717/peerj.6223

16. Dangi, M., Kumari, R., Singh, B. & Chhillar, A. K. Advanced in silico tools for designing of antigenic epitope as potential vaccine candidates against coronavirus. In Bioinformatics: Sequences, Structures, Phylogeny (ed. Shanker, A.) 2018, 329–357

17. Kumar, S., Stecher, G. & Tamura, K. MEGA7: Molecular evolutionary genetics analysis version 7.0 for bigger datasets. Mol. Biol. Evol. 2016, 33(7), 1870–1874. https://doi.org/10.1093/molbev/msw054

18. R. Linding, L.J. Jensen, F. Diella, P. Bork, T.J. Gibson and R.B. Russell Protein disorder prediction: implications for structural proteomics Structure 2003, Vol 11, Issue 11

19. Choi Y, Sims GE, Murphy S, Miller JR, Chan AP. Predicting the Functional Effect of Amino Acid Substitutions and Indels. PLoS ONE 7(10), 2012: e46688 https://doi.org/10.1371/journal.pone.0046688

20. Emidio Capriotti, Piero Fariselli, Rita Casadio I-Mutant2.0: predicting stability changes upon mutation from the protein sequence or structure. Nucleic Acids Research, 2005, Volume 33, Issue suppl_2, W306–W310, https://doi.org/10.1093/nar/gki375

21. Hanka Venselaar, Tim A H Te Beek, Remko K P Kuipers, Maarten L Hekkelman, Gert Vriend Protein structure analysis of mutations causing inheritable diseases. An e-Science approach with life scientist friendly interfaces. BMC Bioinformatics 2010, 11:548. doi: 10.1186/1471-2105-11-548.

22. Gasteiger, E. et al. Protein identifcation and analysis tools on the ExPASy server. In Te Proteomics Protocols Handbook 2005, 571–607, https://doi.org/10.1385/1-59259-890-0:571

23. Cheng, J., Randall, A. Z., Sweredoski, M. J. & Baldi, P. SCRATCH: A protein structure and structural feature prediction server. Nucleic Acids Res. 2005, 33, 72–76. https://doi.org/10.1093/nar/gki396

24. Ashkenazy H, Abadi S, Martz E, et al. ConSurf 2016: an improved methodology to estimate and visualize evolutionary conservation in macromolecules. Nucleic Acids Res. 2016;44(W1):W344–W350. doi:10.1093/nar/gkw408

25. Gaëlle Debret, Arnaud Martel, Philippe Cuniasse RASMOT-3D PRO: a 3D motif search webserver, Nucleic Acids Research, 2009 Volume 37, Issue suppl_2, 1 July 2009, Pages W459–W464, https://doi.org/10.1093/nar/gkp304

26. Pettersen EF, Goddard TD, Huang CC, Couch GS, Greenblatt DM, Meng EC, Ferrin TE. UCSF Chimera—a visualization system for exploratory research and analysis. J Comput Chem 2004;25:1605–1612 https://doi.org/10.1002/jcc.20084

27. Yan Y, Tao H, He J, Huang S-Y. The HDOCK server for integrated protein-protein docking. Nature Protocols, 2020; doi: https://doi.org/10.1038/s41596-020-0312-x.

28. Ankita Barik Mutation Hotspot Prediction, Structural Analysis & Stability Changes of Prion Protein Involved in Causing CJD. Helix 2017, Vol. 8: 1479–1484 DOI 10.29042/2017-1479-1484

29. Yongwook Choi, Agnes P. Chan, PROVEAN web server: a tool to predict the functional effect of amino acid substitutions and indels, Bioinformatics, 2015 Volume 31, Issue 16, Pages 2745–2747, https://doi.org/10.1093/bioinformatics/btv195

30. Linden R The Biological Function of the Prion Protein: A Cell Surface Scaffold of Signaling Modules. Front. Mol. Neurosci. 2017, 10:77. doi: 10.3389/fnmol.2017.00077

31. Hong Ji, Jianfa Wang, Jingru Guo, Yue Li, Shuai Lian, Wenjin Guo, Huanmin Yang, Fanzhi Kong, Li Zhen, Li Guo, Yanzhi Liu, Progress in the biological function of alpha-enolase, Animal Nutrition, 2016, Volume 2, Issue 1, Pages 12–17, https://doi.org/10.1016/j.aninu.2016.02.005.

32. Sofia Khan, Mauno Vihinen Performance of protein stability predictors. Volume 31, Issue 6, Pages 675–684 Wiley InterScience 2010 (www.interscience.wiley.com). DOI 10.1002/humu.21242

33. Mannini, B., Mulvihill, E., Sgromo, C., Cascella, R., Khodarahmi, R., Ramazzotti, M., Dobson, C. M., Cecchi, C., and Chiti, F. Toxicity of Protein Oligomers Is Rationalized by a Function Combining Size and Surface Hydrophobicity. ACS Chem. Biol. (2014), 9 (10), 2309–2317, DOI: 10.1021/cb500505m

34. Kelsey N. Ryan Qixin Zhong Edward A. Foegeding Use of Whey Protein Soluble Aggregates for Thermal Stability—A Hypothesis Paper. CONCISE REVIEWS/HYPOTHESES IN FOOD SCIENCE 2013;Volume 78, Issue 8, R1105–R1115.https://doi.org/10.1111/1750-3841.12207.

35. Droppa-Almeida, D., Franceschi, E. & Padilha, F. F. Immune-informatic analysis and design of peptide vaccine from multi-epitopes against Corynebacterium pseudotuberculosis. Bioinform. Biol. Insights 2018, 12, 1177932218755337. https://doi.org/10.1177/1177932218755337.

36. Shey, R. A. et al. In-silico design of a multi-epitope vaccine candidate against onchocerciasis and related flarial diseases. Sci. Rep 2019. 9(1), 4409. https://doi.org/10.1038/s41598-019-40833-x

37. Meunier, M. et al. Identifcation of novel vaccine candidates against campylobacter through reverse vaccinology. J. Immunol. Res. 2016, 5715790. https://doi.org/10.1155/2016/5715790

38. Hansson, M., Nygren, P. A. & Stahl, S. Design and production of recombinant subunit vaccines. Biotechnol. Appl. Biochem 2000. 32(2), 95–107. https://doi.org/10.1042/ba20000034

39. Capra JA, Singh M. Predicting functionally important residues from sequence conservation. Bioinformatics. 2007;23(15):1875–82. doi: 10.1093/bioinformatics/btm270. Epub 2007 May 22. PMID: 17519246.

40. Albini, A., Di Guardo, G., Noonan, D.M. et al. The SARS-CoV-2 receptor, ACE-2, is expressed on many different cell types: implications for ACE-inhibitor- and angiotensin II receptor blocker-based cardiovascular therapies. Intern Emerg Med. 2020, 15, 759–766. https://doi.org/10.1007/s11739-020-02364-6

41. Lensink, M. F. et al. Blind prediction of homo- and hetero-protein complexes: the CASP13-CAPRI experiment. Proteins, 2019, 87, 1200–1221.

42. Benjamin R Dudenhoeffer, Hans Schneider, Kristian Schweimer, Stefan H Knauer SuhB is an integral part of the ribosomal antitermination complex and interacts with NusA, Nucleic Acids Research, 2019, Volume 47, Issue 12, Pages 6504–6518, https://doi.org/10.1093/nar/gkz442

43. Amar Deep, Prabhakar Tiwari, Sakshi Agarwal, Soni Kaundal, Saqib Kidwai, Ramandeep Singh, Krishan G Thakur Structural, functional and biological insights into the role of *Mycobacterium tuberculosis* VapBC11 toxin–antitoxin system: targeting a tRNase to tackle mycobacterial adaptation, Nucleic Acids Research, 2018, Volume 46, Issue 21, Pages 11639–11655, https://doi.org/10.1093/nar/gky924

44. Sheng-You Huang Search strategies and evaluation in protein–protein docking: principles, advances and challenges, Drug Discovery Today, 2014, Volume 19, Issue 8, Pages 1081–1096, ISSN 1359-6446, https://doi.org/10.1016/j.drudis.2014.02.005.

